# BulkLMM: Real-time genome scans for multiple quantitative traits using linear mixed models

**DOI:** 10.1101/2023.12.20.572698

**Authors:** Zifan Yu, Gregory Farage, Robert W. Williams, Karl W. Broman, Śaunak Sen

## Abstract

Genetic studies often collect data using high-throughput phenotyping. That has led to the need for fast genomewide scans for large number of traits using linear mixed models (LMMs). Computing the scans one by one on each trait is time consuming. We have developed new algorithms for performing genome scans on a large number of quantitative traits using LMMs, BulkLMM, that speeds up the computation by orders of magnitude compared to one trait at a time scans. On a mouse BXD Liver Proteome data with more than 35,000 traits and 7,000 markers, BulkLMM completed in a few seconds. We use vectorized, multi-threaded operations and regularization to improve optimization, and numerical approximations to speed up the computations. Our software implementation in the Julia programming language also provides permutation testing for LMMs and is available at https://github.com/senresearch/BulkLMM.jl.

## Introduction

Genome scans, where genetic variants across the genome are tested for association with traits of interest, are an important tool to discover insights into the etiology of a trait or disease. Recent advancements in high-throughput technologies make it possible to collect large number of traits in a single individual from a single assay. Examples include studies with transcriptomics, metabolomics, microbiome, etc. A common first step in analyzing these data is to compute a genome scan of each trait. This can be a computational challenge since many thousands of traits may be measured. These computational challenges are magnified when linear mixed models (LMMs), the standard approach for genetically structured populations (Li and Zhu, 2013), are used since LMMs are more computationally demanding than linear models. In this work, we tackle the problem of computing genome scans for a large number of quantitative traits using LMMs with the goal of providing runtimes of a few seconds for populations of modest size.

As an example dataset, consider the liver proteome data from the BXD Longevity Study which measured approximately 35K liver proteins on 150 mice from 50 BXD strains (Ashbrook *et al*., 2021). Here, the goal is to map the associations between all 35K liver proteins and approximately 7K genetic markers. The standard approach is to fit the LMM linear mixed model for each protein and marker. This amounts to about 245 million LMM fits. If one is using a LMM genome scan tool such as GEMMA (Zhou and Stephens, 2012), then that program has to be run 35K times. In addition to the repetitive work having to run the program repeatedly, the runtime can be overwhelming even if the tasks were distributed and processed in parallel. When using an interactive web such as GeneNetwork (Sloan *et al*., 2016) speed is key as the user is expecting an answer in less than a minute or even seconds. For interactive analysis, the user may be prepared to sacrifice some accuracy to get a quick overview of the main traits and markers associated with each other; a more accurate and computationally intensive can be done as a follow up. Thus, our goal was to design an algorithm that could complete the analysis of the BXD liver proteome data in a few seconds; we were prepared to make some approximations to accomplish that.

Our implementation, which we call “BulkLMM” (for performing LMMs on a lot of traits “in bulk”), uses ideas for speeding up linear model scans for many traits, combined with techniques for speeding up univariate LMMs, optimization techniques, and efficient implementation using the Julia programming language (Bezanson *et al*., 2017).

The problem of efficiently computing genome scans for a large number of traits using linear models was tackled by Shabalin (2012) who showed that the scans can be greatly speeded up using matrix multiplication instead of performing scans one by one for each marker and trait. The main reason for the speedup is that there are efficient algorithms for matrix multiplication. The tensorQTL (Taylor-Weiner *et al*., 2019) and LiteQTL (Trotter *et al*., 2021) packages used GPUs to speed up the computations further. They use the fact that matrix multiplication uses similar operations with different data, which is an ideal candidate for GPU computation. For computing the scans using LMMs, however, we have to modify the approach used for linear models.

For speeding up the LMM, we use ideas from the FaSTLMM (“Factored Spectrally Transformed Linear Mixed Models”) family of algorithms (Broman *et al*., 2019; Kang *et al*., 2010, 2008; Lippert *et al*., 2011; Zhou and Stephens, 2012). Roughly speaking, FaSTLMM speeds up maximum likelihood estimation by first transforming the data by the spectral decomposition kinship matrix used to express the genetic relatedness of individuals. This effectively transforms the problem into a weighted linear regression problem, which can then be efficiently solved using standard algorithms. To speed the LMMs in bulk further, we use ideas from GridLMM (Runcie and Crawford, 2019) wherein a finite grid of parameter values is considered for optimization. The core idea of BulkLMM is to use highly optimized vectorized and matrix operations whenever possible (as in LiteQTL) and make judicious choices in the FaSTLMM algorithmic pipeline to reduce expensive operations without sacrificing too much accuracy.. In addition to its main functionality designed for fast LMM scans of multiple traits, BulkLMM also provides a fast computation of permutation testing on a single trait (Abney, 2015) and offers features for stabilizing numerical computations.

The remaining article is organized as follows. In Section 2, we outline the modeling framework and general algorithm for fitting the model. In Section 3, we detail our computational methods for speeding up genome scans for multiple traits and give an overview of the methods, including techniques for stabilizing numerical computations. In section 4, we analyze two datasets using our implementation highlighting the runtime performances of our methods in comparison with existing methods. Through our experimentation running our package to perform association mapping on more than 32k expression traits, we demonstrate that BulkLMM has achieved significant runtime improvements over other popular software tools. We end in section 5 by summarizing our conclusions, detailing scenarios suitable for analysis using BulkLMM, and outlining future directions.

## Statistical framework

In this section we outline our statistical approach beginning with a description of the LMM, following with the steps required to fit the LMM, and ending with our approach to permutation testing.

### Linear mixed models (LMMs)

Consider the situation where we have *m* traits and *p* markers measured on *n* individuals. Let *y*_*i*_ denote the *i*-th trait vector (*i* = 1, 2, …, *m*) and *g*_*j*_ (*j* = 1, …, *p*) denote the *j*-th marker coded as allele dosage (or taking values between 0 and 1). Assume the following generative model for *y*_*i*_:

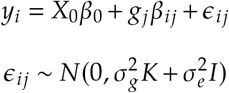

Here, the matrix *X*_0_ contains the covariates that are independent of the tested marker *g*_*j*_, and the vector *β*_0_ contains the corresponding coefficients. We let the marker effect be specifically noted by the indices of trait and marker as *β*_*ij*_. Then, (*X*_0_*β*_0_+ *g*_*j*_*β*_*ij*_) becomes the systematic component of the model.

The random component *ε*_*ij*_ contributes to the variances in the expression trait, which we assume to come from two subsequent variance components: 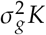 and 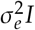. We denote the proportion of total variance explained by genetic variants as 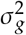 and the remaining unexplained variance as 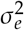. The *n* × *n* matrix *K*, usually referred as the kinship matrix, measures pairwise relatedness identical by descent between each pair of two individuals. Here, we further define the heritability parameter *h*^2^, as 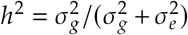, which denotes the ratio of the genetic variance to the total variance. In this way, we may re-parameterize both variance component parameters 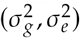 using 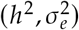, and we emphasize that *h*^2^ is bounded in the interval [0, 1). We will later explain how such reparameterization facilitates our estimation algorithms. For each genome scan, we aim to test the hypothesis of no marker effect (*H*_0_ : *β*_*ij*_ = 0). Essentially, to run genome scans through all pairs of traits and markers, we perform a one-degree-of-freedom test for each pair.

### Fitting the LMM

We are taking a similar approach to the FaST-LMM algorithm. For simplicity of notation, we omit the subscripts and denote simply *y* and *g* as the trait and marker of interest for each test. We define *X* = [*X*_0_, *g*] to be the design matrix of the tested marker and additional covariates *X*_0_ independent to *g*. The fitting of an LMM consists of the following steps.

#### Decorrelation

Given the spectral decomposition of the kinship matrix *K* = *UDU*^*T*^, where the diagonal matrix *D* contains the eigenvalues of *K* on the diagonal and *U* is the matrix with columns of the corresponding eigenvectors, we rotate the original *y* and *X* by (*y*^*^, *X*^*^)= *U*^*T*^ (*y, X*);

After rotation, the transformed data are distributed as

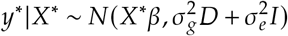

We denote the ratio of the two variance components by *δ*, such that 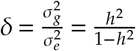. Then, the covariance structure can be written as

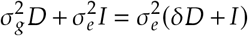

We note that the covariance of the rotated trait is defined by a diagonal matrix with diagonal elements 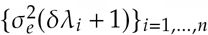 where *λ* _*i*_ is the *i*-th eigenvalue of *K*. Therefore, after rotation, we will have independent observations in the rotated trait, each with the heteroskedastic (unequal) marginal variance defined by the two variance components 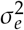, *δ* (in terms of *h*^2^) and a certain eigenvalue *λ* _*i*_. We may then apply the maximum-likelihood principle, or more specifically, the weighted least-square (WLS) approach for estimating the fixed marker effect and parameters of the two variance components.

#### Weighted Least-Squares (WLS)

We write out the log-likelihood function after observing the transformed data (*y*^*^, *X*^*^), as

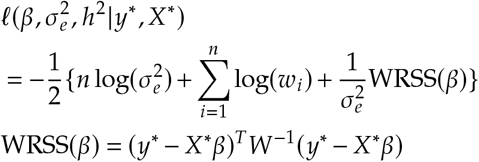

Here, *W* is the diagonal matrix with diagonal elements *w*_*i*_ = *δ λ*_*i*_ + 1, where again 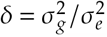, for *i* = 1, …, *n*.

Assuming that the kinship matrix *K* is given and, therefore, its spectral decomposition is known, we notice that the matrix *W* only depends on the unknown parameter *δ*. Given *δ*, we derive the maximum-likelihood estimates of the parameters *β* and 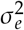 in closed form:

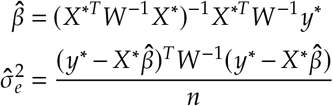

This step is equivalent to estimation by the weighted regression taking each weight as 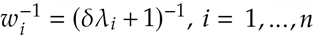.

#### Optimization of h^2^

Plugging the two closed-form solutions for the parameters *β* and σ ^2^, we notice that the loglikelihood function for the data can be seen as a function of only the single parameter *δ*.

In order to better estimate this parameter, we parameterize it using 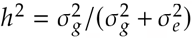, which has physical meaning as the proportion of variance due to genetic variants from the total variance and is bounded in the interval [ 0, 1).

Solving for the estimate of *h*^2^ that maximizes the objective function will finally give us the estimates 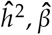, and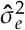 that jointly maximize the likelihood. We applied a one-parameter optimization algorithm Brent’s method (Brent, 1971) to solve for the estimated value of parameter 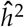.

### Permutation testing

Our approach to permutation testing combines the approach of Abney (2015) with LiteQTL. The essential idea is that given the heritability estimate and, therefore, the weight matrix *W*, we can reweight the observations so that the residuals have zero mean and unit variance. Under the normality assumption, they are also independent under the null. We can permute them several time and reconstruct the trait under the null hypothesis. We can then apply the LiteQTL (matrix multiplication) approach to calculate fit the model under the null.

After being de-correlated and re-weighted,

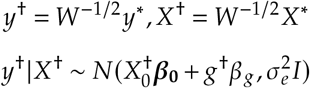

where we used the notation *g* and *β*_*g*_ to denote the tested marker and the corresponding effect. Under the null hypothesis of no marker effect,

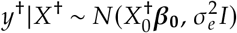

Note that the transformed trait measurements are independent but can have unequal means, as the matrix of control covariates 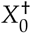 are usually not the identity matrix. Regressing out the covariates gives us independent, identically distributed (i.i.d.) residuals *r*_0_ that since

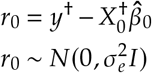

Simply permuting observations in *r*_0_ allows to generate samples that are i.i.d. under the null assumption. Then, following the procedures of permutation test we perform the evaluation scheme previously demonstrated on each of the permuted trait (vector of the same length as *y*) to estimate the fixed marker effect and the resulting testing statistic of LOD. Empirical distribution of the LOD scores from such permutation test framework then allows us to derive estimates of the thresholding values for determine the significance of the marker of each scan.

### Computational methods

In this section, we detail the computational strategies we used to speed up the computation.

#### Heritability estimation precision

The computational complexity of the LMM fitting scheme mainly comes from the estimation of the heritability *h*^2^, which requires solving a one-parameter optimization problem of the objective function on *h*^2^, and the numerical method may be expensive. Once *h*^2^ is estimated, the other two parameters have closed forms and can be easily obtained. For the task of scanning multiple markers, using one linear mixed model at a time for testing each marker, the “Exact” estimation, referred to by other relevant work, assumes that *h*^2^ is independent from one model to another. By “Exact” estimation, We mean that *h*^2^ will be re-estimated for testing each marker. This is a robust but expensive approach, especially when the number of markers is large. As a simple speed-up approach, the “Null” estimation scheme does not re-estimate *h*^2^ at each marker, instead using the approximate value under the null with the baseline (non-marker) covariates and applies the same estimate to test all markers.

In BulkLMM, for each assumption, we have developed scalable algorithms, all of which perform fast even for scanning a large number of traits and markers. We borrow the names “Exact” and “Null” in the names of these developed algorithms to refer to the two assumptions on estimating *h*^2^ each takes.

In the next subsection, we introduce the key computational technique underlying most of the speed-ups in of our proposed algorithms.

### Calculating LOD scores using matrix operations

Let’s assume each trait can be modeled by a simple linear regression with a single covariate; then, based on the fact that the Pearson correlation and the *R*^2^ of testing the single independent variable are equal in this case, we may write the LOD score as

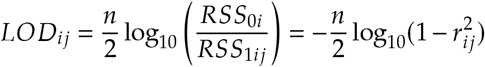

as a function of the correlation coefficient *r*_*ij*_ between each pair of trait *y*_*i*_ and marker *g*_*j*_.

For a set of traits and markers, we can construct the matrices *Y* and *G* with each column in *Y* being a trait and each column in *G* corresponding to the marker to be tested. Then, after standardizing the matrices *Y* and *G* such that each column has zero mean and unit norm, their pairwise correlations can be efficiently computed by a single matrix multiplication (Shabalin, 2012)

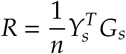

where

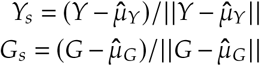

Finally, to convert the pairwise correlation coefficients to LOD scores, we only need to map each element in *R* by the simple one-parameter formula. By this scheme, we do not have to perform a linear regression for each pair of traits and marker, and calculations of LOD scores of all traits and marker pairs can be done efficiently by operations on matrices for which highly optimized implementations are available.

### Accelerating genome scans for multiple traits

To adapt the matrix multiplication technique for bulk LOD score calculations in linear mixed models, we observe that by de-correlating and re-weighting the original data *y* and *X* using a given matrix *W*, we achieve independent transformed data with uniform error variances. This allows us to apply the efficient approach used in simple linear regression. Therefore, the main difficulty lies in how we can reasonably estimate the weight matrix and, consequently, the heritability parameter for different traits.

For performing scans on a single trait, this matrix multiplication scheme can be applied by replacing the matrix *ϒ*^†^ with a single trait column matrix *y*^†^ and using the full genotypes at all tested markers to construct matrix *X*^†^. Then, by matrix multiplication and mapping the pairwise correlations, we can efficiently compute the LOD score between the single trait and every marker. This idea has led to our first algorithm by naively scanning one trait at a time with the exact estimate of heritability for each trait.

For convenience in our further demonstration of the various algorithms we developed, we will use 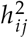 with the two subscripts (*i, j*) to denote the heritability parameter for a particular trait *y*_*i*_ and marker *g*_*j*_, where *i* = 1, …, *m* (total number of traits) and *j* = 1, …, *p* (total number of markers). Specifically, we use 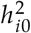 with *j* = 0 to denote the heritability under the null model when there is no marker effect for each trait *y*_*i*_. To also differentiate between the use of the term “Exact” by other relevant work, which indicates that the heritability for each marker will be estimated independently, and our use of the term “Exact” in the method’s name to emphasize that the estimated value of *h*^2^ is from optimizing the actual objective function using numerical methods rather than grid-search, we will use the different term “Alt” (versue “Null”) to have the same meaning as the “Exact” as the previous work referred to.

#### Based on exact estimation of 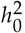

Assuming 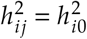 for all *j*, our naive approach to extend the use of the matrix multiplication strategy to our linear mixed model case is to construct the matrix *ϒ* ^†^ in the scheme using one trait *y* at a time while constructing the matrix *X*^†^ using all genome markers, which is outlined as follows:

##### Algorithm 1

Bulkscan: Null-Exact

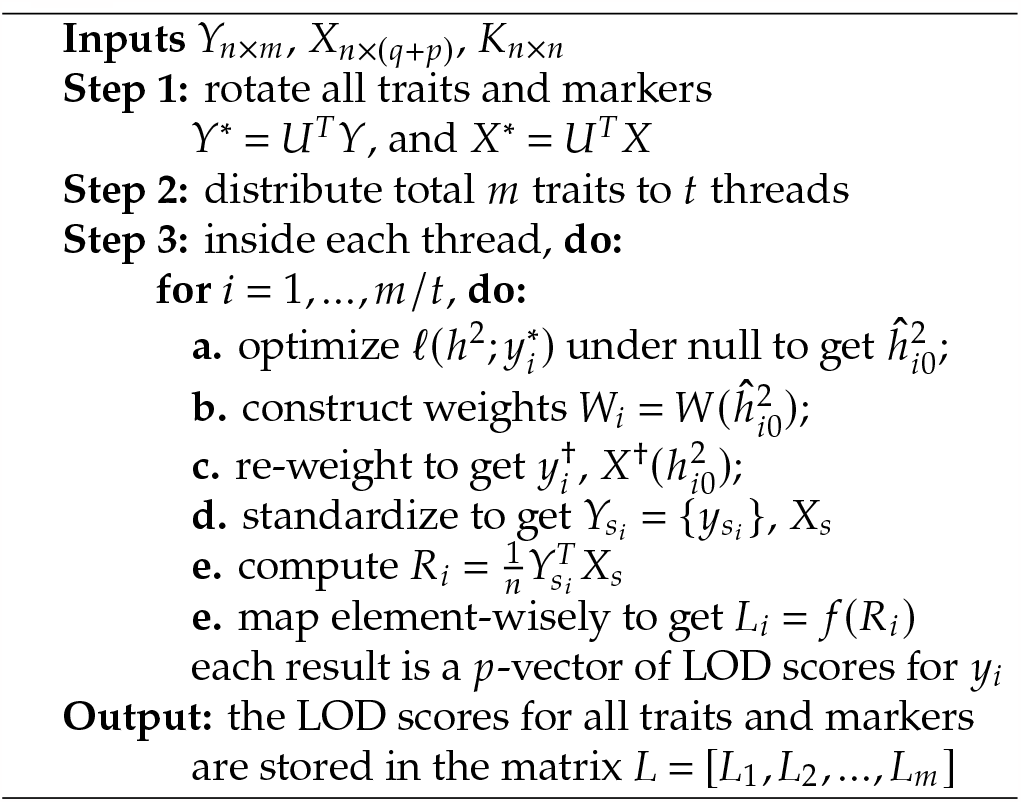

For further speed-ups over our first naive method, it is tempting to think of approaches to construct the matrix *ϒ* ^†^ using not only one but ideally a fair portion of the total number of traits to be tested, then the runtime is expected to be reduced as a fraction of the runtime of the first algorithm. While such an approach to group up traits may not be feasible by their exact estimates of heritability since parameters from optimizing the objective function independently are very unlikely to be exactly the same, some approaches by relaxing the precision from exact estimation are promising to find common estimates for multiple traits, enabling grouping of those traits for applying one matrix multiplication for bulk calculation of their LOD scores. This approximation idea has motivated us to further improve the execution time of our algorithms, and through our experiments, for most cases, estimating the heritability parameter up to some levels of precision can be sufficient to generate results that are reliably accurate.

#### Based on grid-approximation estimation of 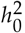

The seond algorithm we propose, named, *Bulkscan-Null-Grid* makes additional relaxation on the accuracy required for the results by estimating the heritability of each trait approximately on a grid of finite candidate values (Runcie and Crawford, 2019). In such a manner, we will have multiple traits with the same heritability estimates. Then, the matrix multiplication approach for computing the LOD scores for traits modeled by linear mixed models can be extended to testing multiple traits instead of one, as our proposed algorithm “Bulkscan-Null-Exact” does. The weighted likelihood function values for all traits are computed under different weights each depending on a candidate *h*^2^. The final *h*^2^ estimate for each trait is determined as the candidate *h*^2^ value that yields the optimal value of the objective function. The next step is to create batches for traits sharing the same heritability estimate from the grid-search step. Within each batch, the traits are used to construct the matrix of responses *ϒ* ^†^ and to perform matrix multiplication as demonstrated to compute the LOD scores for those traits.

##### Algorithm 2

Bulkscan: Null-Grid

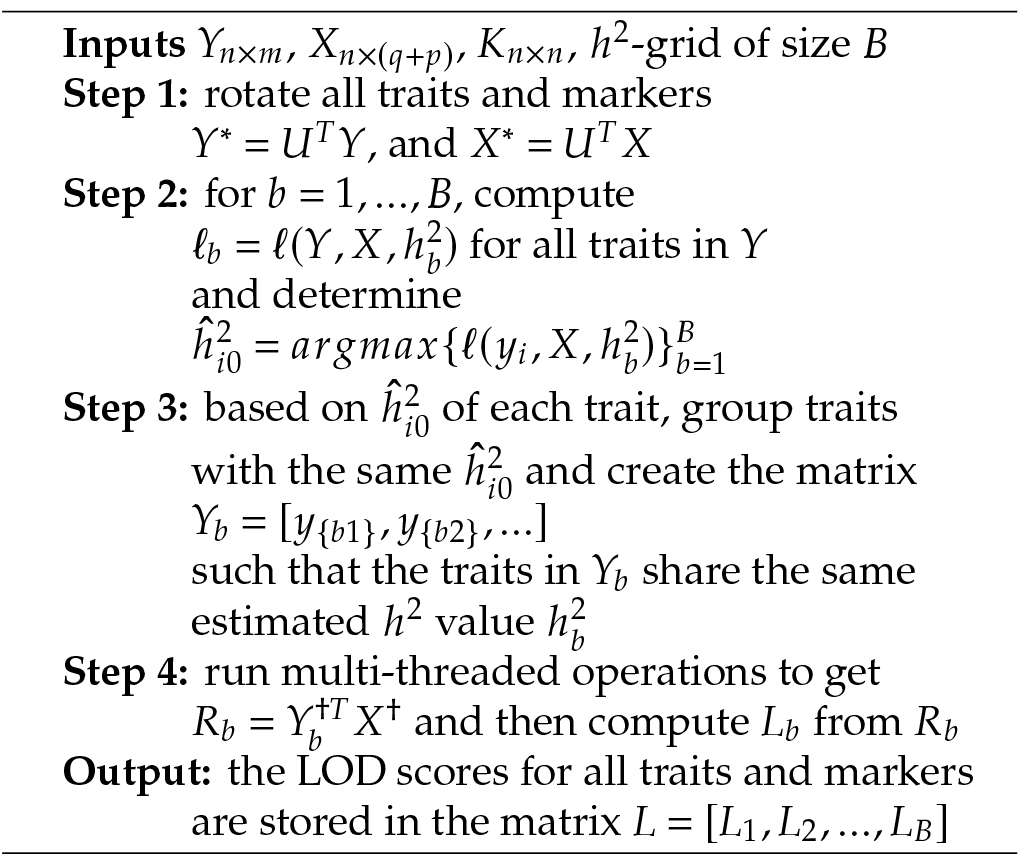

For demonstration of the algorithm, we use the notations 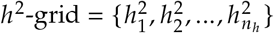 to denote input grid of *n*_*h*_ possible values and let 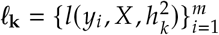 be the values of the objective function for each trait evaluated at a given 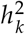. The overview of the algorithm is as follows.

In Algorithm 2, it is important to note that each *h*^2^ value specifies a weight matrix *W*. Consequently, the step of calculating log-likelihood function values for a fixed *h*^2^ can be efficiently executed for all traits through a multivariate weighted regression, using *ϒ* as a response matrix with its columns representing the traits. Then, the *𝓁* _**k**_ of the log-likelihood values for all *m* traits can be seen as a row vector of length *m*. After computing the objective function values for all values of 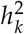, we can stack the row vector *𝓁* _**k**_’s to form a matrix. Therefore, finding the optimal function value and the corresponding parameter value *h*^2^ for each trait is done by finding the maximum value in each column of the resulting matrix of log-likelihood values.

#### Based on grid-approximation estimation of 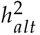

The third method we propose, named *Bulkscan-Alt-Grid*, combines the ideas of the grid-search approach for estimating the heritability and the matrix multiplication approach for efficiently computing LOD scores. It computes the LOD score based on heritability estimated independently from each marker tested for each trait.

A key observation is that for each test of a trait *y*_*i*_ and the design matrix *X*_*j*_ containing the tested marker *g*_*j*_, from the formula of the LOD score that

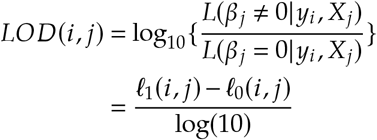

we can recover

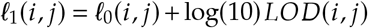

Therefore, under the linear mixed model scheme, for a given *h*^2^ value, we can first apply the matrix multiplication to compute the pseudo-*LOD* (*i, j*) for all *i* and *j*. Then since 𝓁 _0_ (*i, j*) can be calculated easily for all *i* and *j* (essentially because the null log-likelihood does not depend on the specific marker *g*_*j*_) also by matrix operations, we can recover the alternative model log-likelihood under a *h*^2^ for all *i* and *j* from the above derivation. Specifically, these alternative model log-likelihood values will be stored in a matrix of dimension *p* × *m* for each 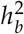 in the grid. By optimizing element-wisely the 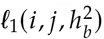, we can get the estimated alternative model log-likelihood for each trait *y*_*i*_ and marker *g*_*j*_,under the optimal value of heritability for each alternative model containing each specific marker *g*_*j*_. Finally, the true LOD score evaluated under the optimal 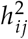 and

#### Algorithm 3

Bulkscan: Alt-Grid

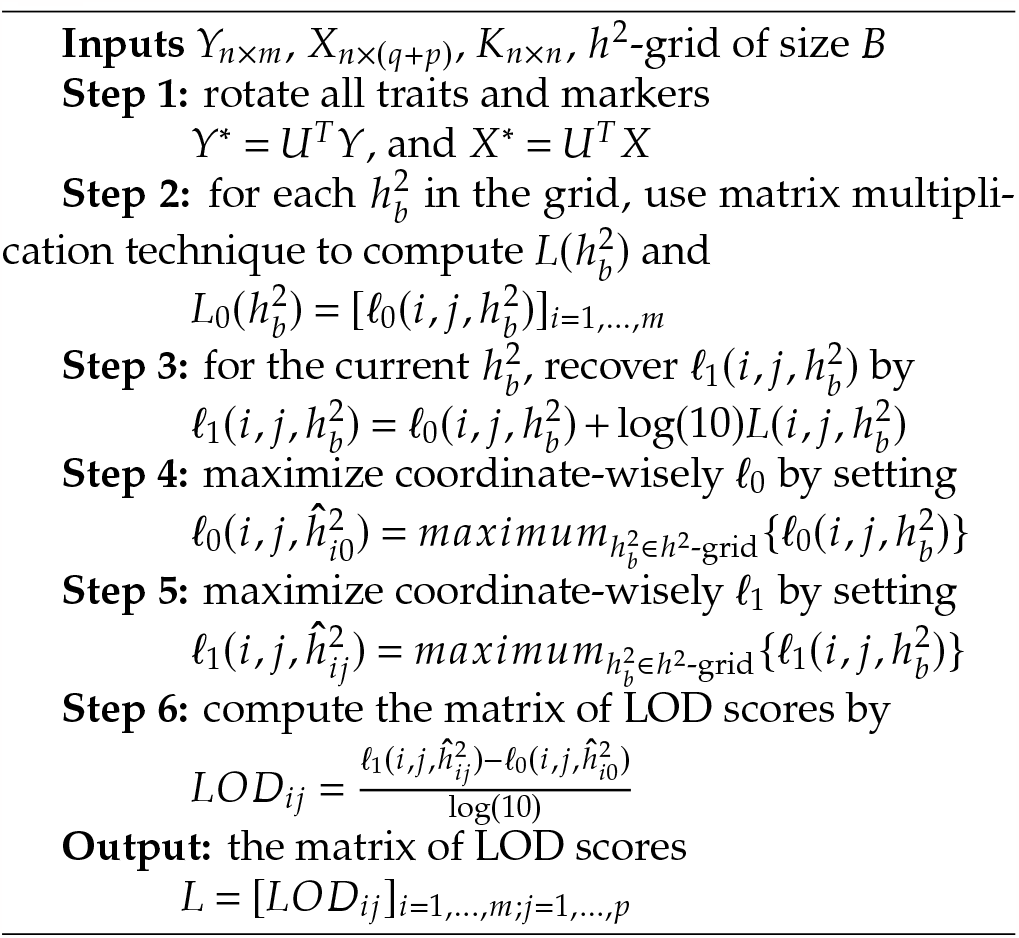

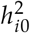 for each *i* and *j* can be calculated based on

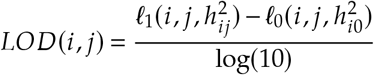

where 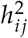 and 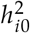 are each the optimal heritability estimated from the alternative model containing the marker *g*_*j*_ and the null model, respectively.

### Numerical stabilizing techniques

When performing a large number of genome scans in parallel, the chance of encountering unlikely situations is increased. Therefore, in addition to speed, we have to also pay attention to numerical stability because otherwise, a whole batch of computations can fail because of one unusual trait or marker. We first discuss boundary avoidance, which is focused on techniques to avoid situations when the heritability is exactly 1. We follow with an improvement to heritability estimation using Brent’s method by sudividing the unit interval into subintervals to avoid multiple local maxima.

#### Boundary Avoidance

Notice that if the estimated 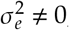, the objective function maximized at the closed-form, maximum-likelihood estimators 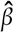and 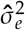for optimization on *h*^2^ can be written as

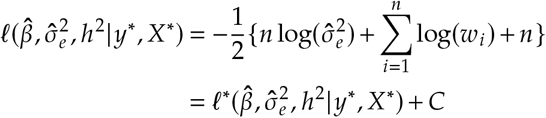

where *C* = −*n*/2 is a constant term.

Let us look carefully log-likelihood function as the heritability estimate *h*^2^ approaches 1. We write the above function as

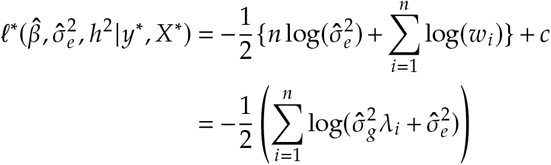

As 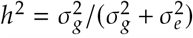 approaches 1, 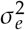 approaches 0. The log likelihood will blow up to infinity if there is at least one *λ* _*i*_ ⋍ 0. That is possible when the kinship matrix is not full rank, for example when two or more individuals have the same genotype at all markers.

To correct the numerical issue of heritability being estimated at 1, we take a Bayesian maximum a posteriori (MAP) approach for estimating the residual variances 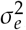, by imposing a prior on 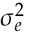 during its estimation (Galindo Garre and Vermunt, 2006). The prior distribution represents our prior belief that the residual variances are very unlikely to be 0. Specifically, we implement the prior distribution of a Scaled-Inverse-*χ* ^2^ on the 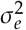, with scale parameter 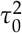 and *χ* ^2^ degrees of freedom *v*_0_, and a support of (0, ∞). Therefore, rather than estimating 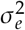 by maximizing the log-likelihood function, we estimate by maximizing the log posterior distribution, where the posterior is proportional to the product of the prior and the likelihood function of the data:

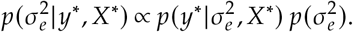

We take this prior choice mainly for taking the computational advantage of the Inverse-*χ*^2^-Normal conjugacy, for easily evaluating the a posteriori without the need to evaluate the integral for marginal distribution of the data (Gelman *et al*., 2013). Finally, for estimating the heritability *h*^2^, we plug in the MAP estimates of 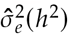 and 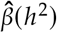 to the log-posterior as the final objective function and apply numerical optimization methods, similarly as the approach of no added prior.

#### Sub-regional numerical optimization using the Brent’s method

When we estimate the variance components of the LMM by optimizing the heritability we apply Brent’s method over 0, 1. The `Optim.jl` package in Julia provides an implementation of this method. However, Brent’s method is sensitive to the initial guess as well as the shape of the objective function and can produce incorrect results if the objective function has more than one local minimum over the optimization interval.

To mitigate these issues, we provide an option to sub-divide the whole optimization region and applying Brent’s method to each sub-interval. The final result is determined by comparing the objective function at the sub-interval roots. The number of sub-divisions is given by the user; more subintervals give greater accuracy at the price of lower speed. In some cases, narrowing down the search space might lead to a better convergence rate for Brent’s method. As the optimization in each sub-interval is independent, even faster computational speed can be achieved if these operations are parallelized.

**Fig. 1.**
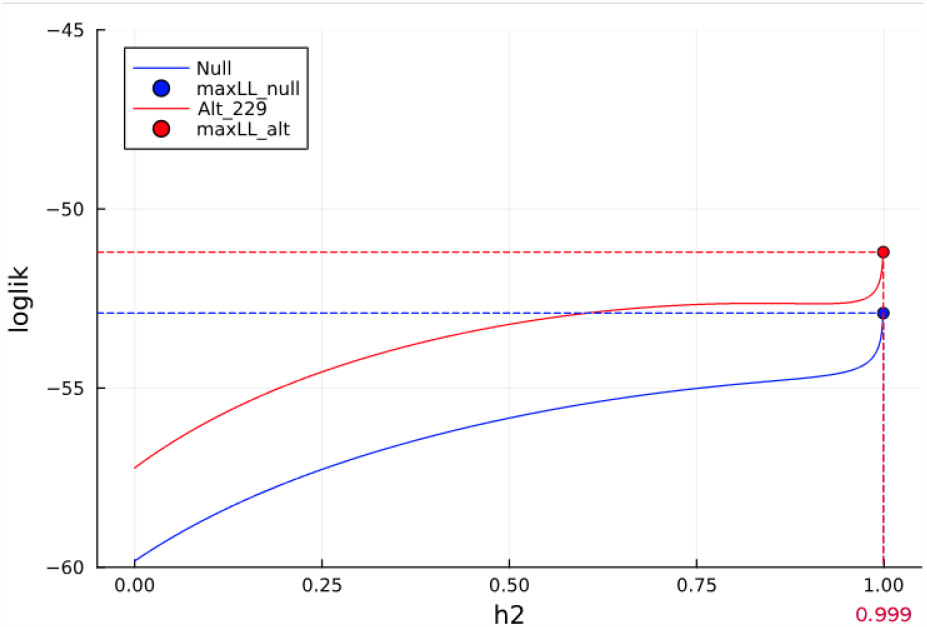
Estimation of heritability by maximum likelihood. Demonstration of this issue is reproducible using the sample BXD Spleen data. We show the curves of the objective function (log-likelihood) under the null model and under the alternative model testing the 229th marker, with each curve consisting of 1000 values under a *h*^2^ value 0.0, 0.001, …, 0.999. Without correction, both objective function values blow up near heritability of 1.

**Fig. 2.**
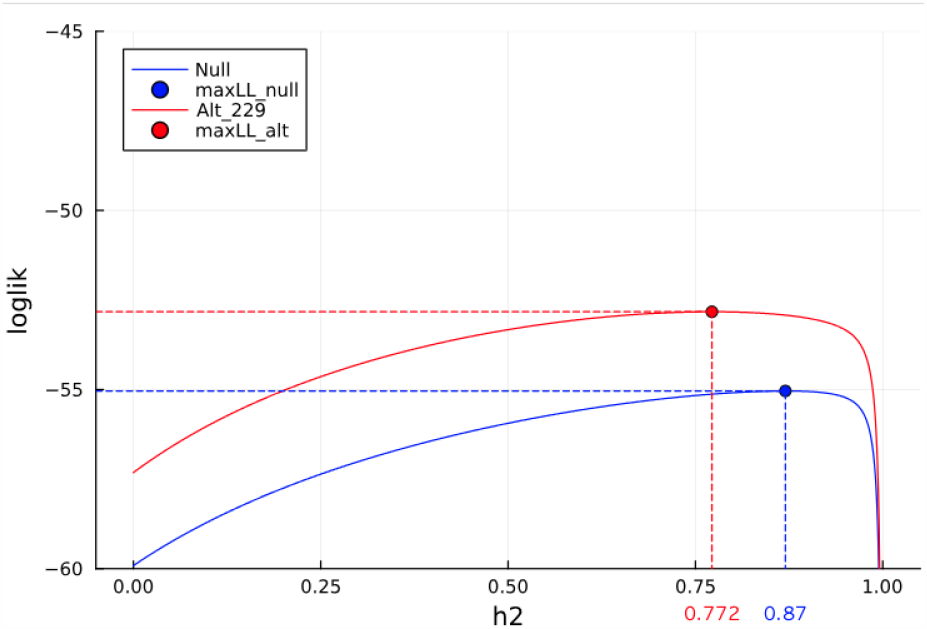
Estimation of heritability by maximize-a-posteriori after imposing the prior Scaled-Inv-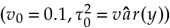. We show the curves of the objective function (log-posterior) under the null model and under the alternative model testing the 229th marker, with each curve consisting of 1000 values under a *h*^2^ value 0.0, 0.001, …, 0.999. With boundary avoidance correction, both objective functions become concave and estimation produces more reliable heritability estimates.

## Results

In order to provide the future users of BulkLMM a comprehensive view of its performances under various scenarios, depending on the sizes of the input data, as well as the options for model evaluation requested by the user, we used BulkLMM to perform two analyses - one on the BXD mice liver proteome traits and the other on the heterogenous stock (HS) rats prelimbic cortex transcriptome. We describe the runtime performance of our algorithms and follow it by an assessment of the accuracy of the algorithms. The goal is to assess the tradeoff between speed and accuracy, so that the user can make informed choices.

### Datasets

The BXD mice liver proteome data consists of individual-level measurements on a total of 32445 liver proteins. The 248 individual samples came from 50 BXD strains genotyped at 7321 markers. This data has a modest sample size with population structure so that the genetic analysis should be done using LMMs.

The second dataset is from 80 heterogeneous stock (HS) rats whose prelimbic cortex transcriptome was measured using 18,416 features. These rats were genotyped at 117,618 markers making it a larger dataset in terms of number of markers.

### Runtime performance

To assess runtime performance of BulkLMM under we executed each BulkLMM method as well as the univariate LMM genome scan in GEMMA on overall *m* (32K for the BXD data and 18K for the HS data) traits and recorded their runtimes. For BulkLMM, we used a 24-threaded Julia session on the most recent stable release of Julia, version 1.9.2, where the optimization of *h*^2^ was based on REML and on a grid of 0.1 *h*^2^-stepsize. Since the method in GEMMA for univariate linear mixed modeling takes only one trait at a time, we iteratively ran GEMMA on 1000 randomly selected traits and approximated the runtimes for processing all *m* traits by the execution times for 1000 traits times *m* 1000. Such extrapolation is reasonable since running GEMMA iteratively for GWASs on multiple traits has runtimes that are approximately linear in the number of traits. A summary of the approximated run-time of each method is shown in Table 1.

**Table 1.** Runtime performances by BulkLMM methods and GEMMA on two datasets. For our two grid-search based methods, namely “Null-Grid” and “Alt-Grid”, the execution times were from running these methods on a coarse grid (0.1) of *h*^2^ (i.e. a grid of 10 values from 0.0 to 0.9 with the stepsize of 0.1). We observed the fastest computational speed on the BulkLMM-null-grid approach; the BulkLMM-null-exact and BulkLMM-alt-grid come next (depending on the dataset); and GEMMA is much slower.

| Data | Method | Runtime |
| --- | --- | --- |
| BXD Liver Proteome | BulkLMM-null-exact | 100s |
|  | BulkLMM-null-grid | 4.8s |
|  | BulkLMM-alt-grid | 52s |
|  | GEMMA (alt-exact) | 450s |
| HS Prefrontal Cortex | BulkLMM-null-exact | 440s |
|  | BulkLMM-null-grid | 14s |
|  | BulkLMM-alt-grid | 570s |
|  | GEMMA (alt-exact) | 220,000 s |

### Numerical accuracy compared to GEMMA

We compared the numerical accuracy of our methods to that from GEMMA, which optimizes the heritability at each marker.

We verified that BulkLMM would generate reliable results by comparing the results from 1000 randomly selected traits in the BXD individual liver proteome data. We compared the results from both methods on the −log_10_(*p*) scale. As a summary, we report the sample mean of the element-wise absolute difference over the total of number of traits (1000) and markers (7321) in Table 2. We see that for “alt-grid” approach with a fine (0.01) grid is almost the same as GEMMA, and that “null-grid” method on a coarse (0.1) grid is the fastest, but the greatest approximation error. This is reinforced in Figure 3 which plots the GEMMA output with the output from the “null-grid” coarse grid, and “alt-grid” fine grid approaches.

**Table 2.** BulkLMM runtimes and accuracy (MAD, mean absolute difference) for specific choices of methods and sizes of *h*^2^ -grid. We use the number in the parenthesis after the name of the method to denote the stepsize of *h*^2^ -grid. The runtimes use 1000 randomly chosen BXD dataset traits, and GEMMA is used for accuracy baseline. We can see that the null-grid approach with a coarse grid (0.1) is the fastest, while still maintaining reasonable accuracy. The alt-grid method with a coarse grid (0.1) is faster than the null-grid method with a fine grid (0.01), while being about as accurate. The alt-grid method with a fine grid (0.01) is almost indistinguishable from GEMMA and is 10 times slower than the alt-grid method with a coarse grid.

| BulkLMM method | MAD | Runtime |
| --- | --- | --- |
| null-exact | 0.0095 | 5.1s |
| null-grid (0.1) | 0.018 | 0.38s |
| null-grid (0.05) | 0.012 | 0.64s |
| null-grid (0.01) | 0.010 | 2.5s |
| alt-grid (0.01) | 0.011 | 1.4s |
| alt-grid (0.05) | 0.0038 | 2.8s |
| alt-grid (0.01) | 0.00097 | 14s |

**Fig. 3.**
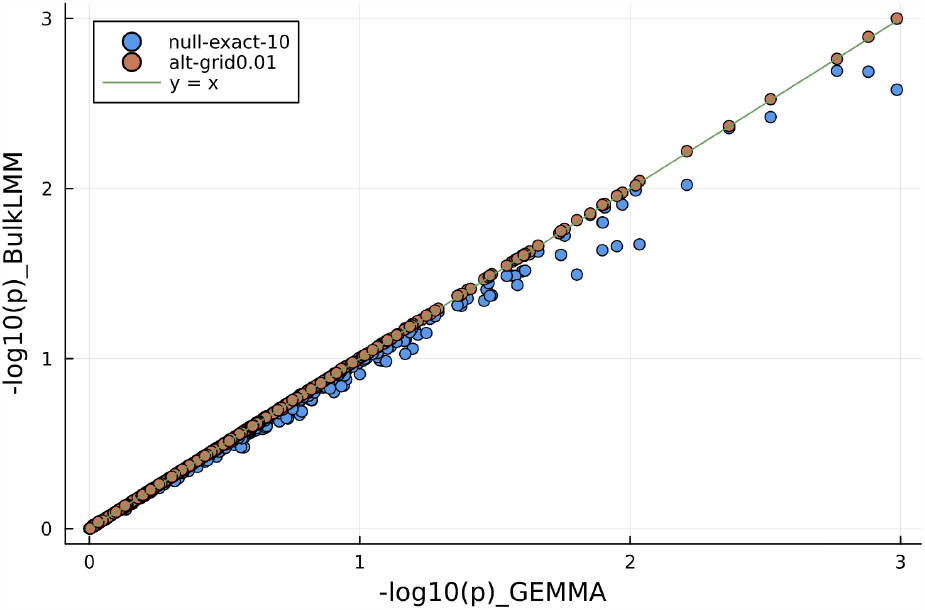
P-values from running “null-exact” and “alt-grid” with a *h*^2^ -grid of 100 values in comparison with GEMMA on the BXD data. 1000 randomly picked p-values were plotted. The closer the points to the solid line suggests better accuracy as to reproduce GEMMA results. In Fig.2, we notice that by using a grid of 100 values representing a resolution of 0.01, our “alt-grid” results are much closer to GEMMA results.

### Runtime and accuracy as a function of *h*^2^-grid size

One of our approaches to speed the LMMs in bulk is to perform a grid search across a discrete set of points in the interval of 0, 1. The performance of our two methods based on grid-search depend on the *h*^2^-grid resolution. For our fastest algorithm, the “null-grid”, which is benefited from and therefore has runtimes affected by such shortcut the most, we ran the method and recorded the execution times under *h*^2^-grid of different step-sizes from 0.01 to 0.1 (corresponding to sizes of *h*^2^-grid from 100 to 10), for performing a scan over all 32k traits of the BXD liver proteome data. We plotted the runtime curve, as well as the curve of the mean deviation of LOD scores compared to LOD scores from the null-exact method, as functions of the *h*^2^-grid size, shown in Figure 4.

**Fig. 4.**
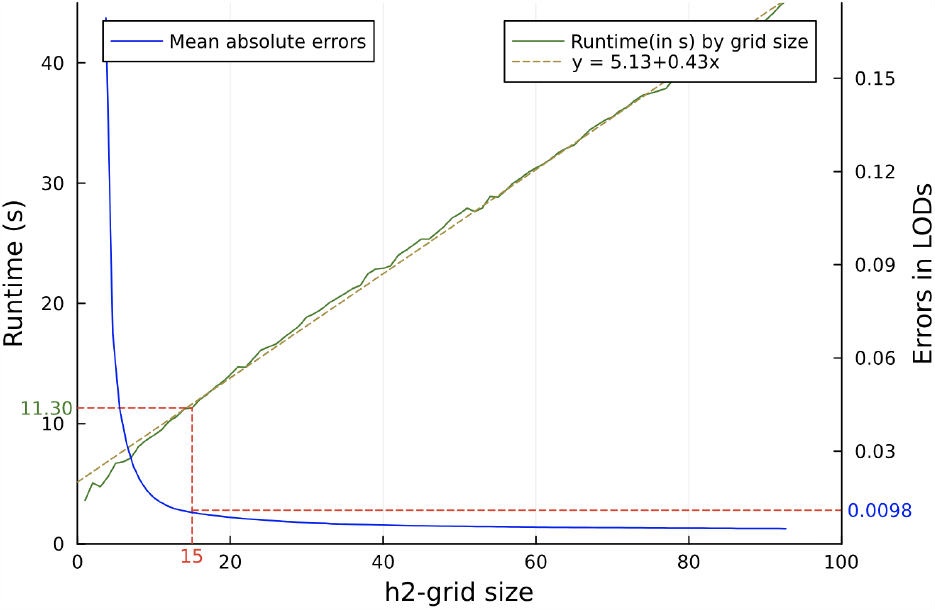
Line plots showing run times and errors from running “null-grid” by different sizes of *h*^2^ -grid, from 1 to 100 values. We observe that runtime is almost linear and error decrease more rapidly for low *h*^2^ -grid sizes. This suggests that while runtime of the null-grid method increases linearly with the sizes of the *h*^2^ -grid, the errors decreases sharply when increasing the sizes from low *h*^2^ -grid sizes, and the marginal gain in accuracy by increasing *h*^2^ -grid size will continue to decrease until no significant marginal impact.

We also compared the accuracy of the “alt-grid” method as a function of grid size, and find that while both are quite accurate, a fine (0.01) grid gives results almost identical to GEMMA (Figure 5).

**Fig. 5.**
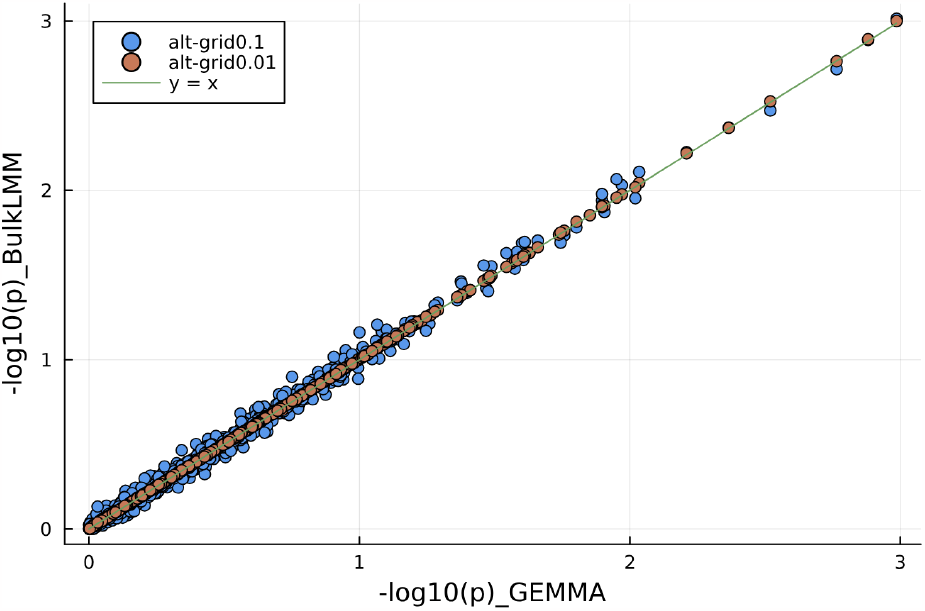
P-values from running “alt-grid” with two grids of 100 and 10 values in comparison with GEMMA on the BXD data. 1000 randomly picked p-values were plotted. The closer the points to the solid line suggests better accuracy as to reproduce GEMMA results. In Fig.3 we further notice that by using a grid of 10 values representing resolution of 0.1, our “alt-grid” results are still not so different as compare to GEMMA.

### Adjustment of sample relatedness by using LMMs

The most fundamental reason for favoring the linear mixed models over linear models for GWAS is to control for the genetic relatedness among the sample individuals. We compared the 1000 randomly selected results in the format of −log_10_ (*p*), from running BulkLMM “null-grid” and “alt-grid”, each with a *h*^2^-grid of stepsize 0.01 with the results computed from simply linear models. Figure 6 plots the comparison of these results.

**Fig. 6.**
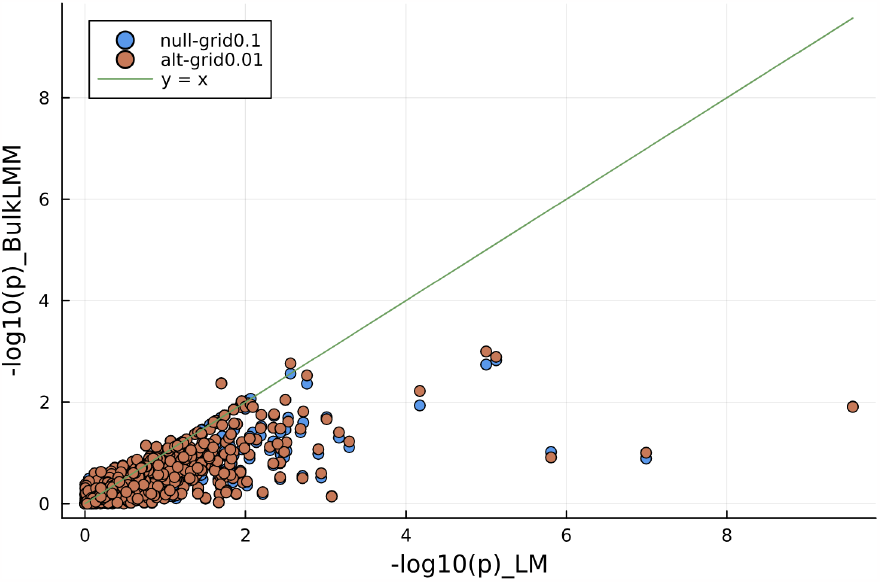
Comparison of results on the individual-level on BXD liver proteome by linear mixed models and by linear models. For linear mixed modeling, results from running “Null-Grid” and “Alt-Grid” with 10 and 100 *h*^2^ values respectively are shown. These results indicate that the −log_10_( *p*)from LMMs are almost always lower than those from the corresponding linear models, suggesting a lower effective sample size due to genetic relatedness.

## Discussion

Our BulkLMM package is able to perform genome scans for thousands of traits in moderately-sized populations in a few seconds (5 seconds for the BXD data, 14 seconds for the HS data). These represent speedups of 94 times and 16,000 times respectively compared to running GEMMA one trait at a time. Running GEMMA one trait at a time scales linearly with the number of traits, and may be infeasible for some datasets. We believe our approach makes these datasets tractable and opens up the possibility of interactive analyses in real time for genome scans for high throughput traits.

### Trade-off between runtime and precision

We were able to achieve our fastest speeds with the “null-grid” method which uses two approximations: (a) it estimates the heritability under the null only and (b) it considers a grid of heritabilities. The method then groups the traits by the best heritability on the grid, and then uses matrix multiplication to efficiently calculate the LOD scores. This method was the least accurate of the three algorithms, “null-exact”, “null-grid” and “alt-grid”, for fitting the LMMs. Our results show precision of the results given by these algorithms as well as the their runtime performances reflecting the slightly different approaches each taken for estimating the heritability. Methods that allow for greater slack in heritability estimation therefore have faster speeds. We let the user choose the approach that fits their needs. For a quick overview, the “null-grid” method is the best, but if greater accuracy is needed, the “alt-grid” method on a finer grid is recommended. Since the speed scales linearly by the number of grid points, we suggest using a coarser grid first before using finer grids.

### Performance characteristics

As for the accuracy of BulkLMM, all of our methods generate reliable results, reflected by the diminutive differences to the results from GEMMA. Based on the results of eQTL analyses of randomly selected 1000 BXD liver proteome traits using BulkLMM and GEMMA, we show that the mean absolute difference in LOD scores is less than 0.02 for our least-accurate but fastest method, and for our most-accurate method which does the exact-LMM similar to GEMMA but taking a grid-search approximating approach, the difference is less than 0.001.

### Impact of data size

Our key computational technique for speeding up genome scans for a large number of traits is to convert the iterative processes to a set of equivalent operations on large matrices, with the sizes of the matrices depending on the number of individuals, phenotypes and genotyped markers. The HS data had a smaller number of individuals and traits, but a much larger number of markers than the BXD dataset. The runtimes for the HS data were much longer, but there is not a simple relationship in the runtimes. In general, we expect the runtimes to also depends on the architecture of the machine (number of threads/cores, CPU speed, bus speeds, and available RAM). For datasets with a very large number of markers, it would be preferable to split up the computation by chromosome or smaller subsets to make the data fit in memory.

### Additional features of the implementation

Although we are focussed on performing LMM genome scans in bulk, we provide some additional features that many users might find useful. First, we provide a permutation testing for a single trait which utilizes the key matrix multiplication approach for efficiently calculating the LOD scores for permuted copies and can perform in real-time. Second, if the residual variances that may not be equal for all individuals, we allow the user to specify a weighting scheme for a weighted LMM. This feature is useful if we have unequal replicates per recombinant inbred line, and we wish to use the strain mean as the trait value. Finally, we provide the option to provide a prior for the residual variance. While our motivation was boundary avoidance, this feature can also be used when summary data is used for the genome scans. We will expand on this topic in a future publication.

### Potential limitations

While we have succeeded in speeding up the process of computing LMM genomne scans in bulk, we made a number of design choices that have consequences.

#### No missing data

The most important practical limitation is that we assume that our phenotype and genotype inputs have no misssing data. The user has to either impute or remove individuals/markers with missing data. Genotypes are routinely imputed and should not present a major obstacle. Imputation of the traits may be more challenging, but with high-quality data this should not be a major limitation.

#### One degree of freedom tests

Our speedups rely on matrix multiplication, and that approach assumes that we are interested in one-degree of freedom tests for genomewide scans. For most GWAS panels and recombinant inbred panels such as the BXD family that is not a problem. Additional work is needed for situations where two or more degree of freedom tests are needed.

#### Single kinship matrix for measuring relatedness

Our LMM framework assumes that genetic relatedness can be adequately adjusted using a single kinship matrix. In some situations (such as when a dominance kinship matrix is desired), that may not be adequate, and additional work would be needed to handle multiple kinship matrices. *Grid search*. The precision of our grid-search methods depend on the shape of objective function as a function of the heritability. If the curvature of the actual function is large near the maximum, then our grid approximation may not fare well. Empirically, this can be tested by using a finer grid and comparing the results. If there is a big change, the finer grid should be used.

#### Variance independent of mean assumption

Our implementation is designed for traits whose variance in idependent of the mean. For count data or binary data, alternative approaches need to be devised. Finally our framework is designed for multiple independent quantitative traits. In some situations, a multivariate linear mixed model may be more suitable (Kim *et al*., 2020).

### Implementation in Julia language

We implemented our software in the Julia programming language (Bezanson *et al*., 2017) which has computational speed comparable to some lower-level languages such as C, C++, but has clean syntax similar to high-level languages such as Python and R. Our implementation used Julia’s support for multithreading. Further speedups may be possible with the use of GPUs and may be a topic of future investigation.

## Data availability

For the two datasets used during our experimentation, the BXD individual liver proteome and the HS rats mRNA data, both are open to public access and are accessible from the GeneNetwork database at https://genenetwork.org/. The BXD liver proteome data of BXD Longevity Study can be obtained by using the file-name *EPFL/ETHZ BXD Liver Proteome CD-HFD (Nov19)* or the accession code: GN886. The Prelimbic Cortex mRNA data of HS-Palmer Rats are accessible using the following query in GN:

- Species: Rat
- Group: NIH Heterogeneous Stock (Palmer)
- Type: Prelimbic Cortex mRNA
- Dataset: HSNIH-Palmer Prelimbic Cortex RNA-Seq (Aug18)
- Get Any: *

